## Supplementary Figures for "A Remarkable Genetic Shift in a Transmitted/Founder Virus Broadens Antibody Responses Against HIV-1"

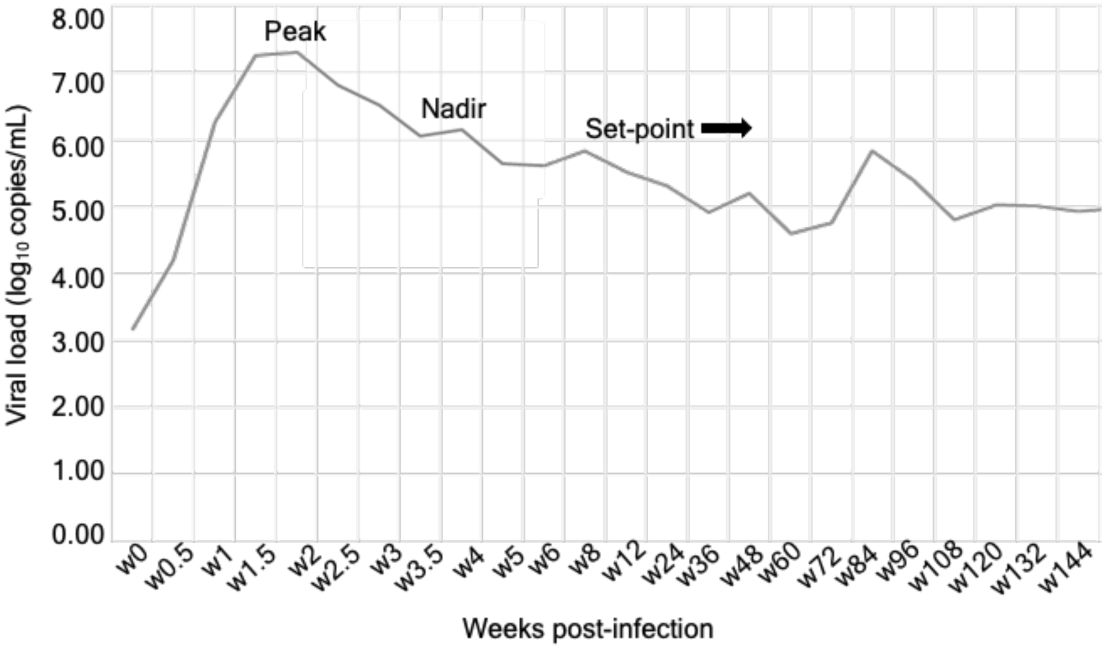


**Fig. S1. Longitudinal viral load analysis over a course of infection in the participant 7.** Viral load values ($log_{10}$) are plotted on y-axis, against the number of weeks since the first HIV-positive reaction with nucleic acid test on the x-axis. A typical pattern of curve with a peak, nadir, and set point in viral load was observed in this patient representing early captured infection. The set point is the viral load of a person infected with HIV, which stabilizes after a period of acute HIV infection.

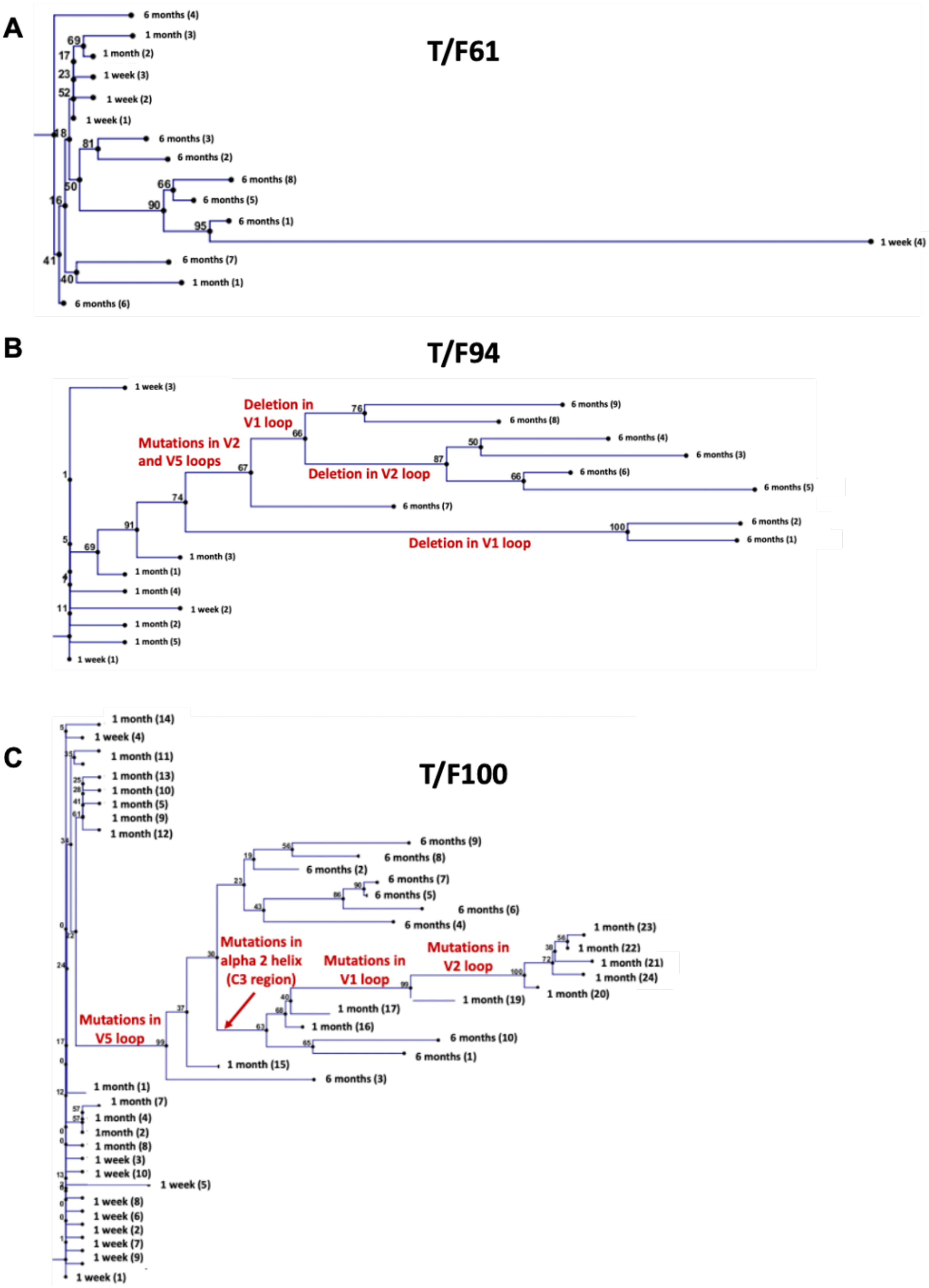
A
T/F61
6 months (4)
1 month (3)
1 month (2)
1 week (3)
1 week (2)
1 week (1)
6 months (3)
6 months (2)
6 months (8)
6 months (5)
6 months (1)
1 week (4)
6 months (7)
1 month (1)
6 months (6)
B
T/F94
1 week (3)
Deletion in V1 loop
Mutations in V2 and V5 loops
Deletion in V2 loop
Deletion in V1 loop
6 months (9)
6 months (8)
6 months (4)
6 months (3)
6 months (6)
6 months (5)
6 months (7)
6 months (2)
6 months (1)
1 month (3)
1 month (1)
1 month (4)
1 week (2)
1 month (2)
1 month (5)
1 week (1)
C
T/F100
1 month (14)
1 week (4)
1 month (11)
1 month (13)
1 month (10)
1 month (5)
1 month (9)
1 month (12)
6 months (9)
6 months (8)
6 months (2)
6 months (7)
6 months (5)
6 months (6)
6 months (4)
Mutations in alpha 2 helix (C3 region)
Mutations in V1 loop
Mutations in V2 loop
Mutations in V5 loop
1 month (23)
1 month (22)
1 month (21)
1 month (24)
1 month (20)
1 month (19)
1 month (17)
1 month (16)
6 months (10)
6 months (1)
1 month (15)
6 months (3)
1 month (1)
1 month (7)
1 month (4)
1month (2)
1 month (8)
1 week (3)
1 week (10)
1 week (5)
1 week (8)
1 week (6)
1 week (2)
1 week (7)
1 week (9)
1 week (1)

**Fig. S2. Phylogenetic trees showing evolution of transmitted/founder viruses.** (A-C) Phylogenetic trees depicting the divergence of T/F61 (A), T/F94 (B) and T/F100 (C) viruses from the time of infection until 6-month post-infection. The trees are constructed with T/F virus sequence as root using neighbor-end joining method in CLC Mains Workbench with parameters as described in Materials and Methods. Dominant mutations resulting in divergence of the viruses are labeled on the branches of the tree. T/F61 viruses lacked any predominant mutations and had conserved V1V2 region.

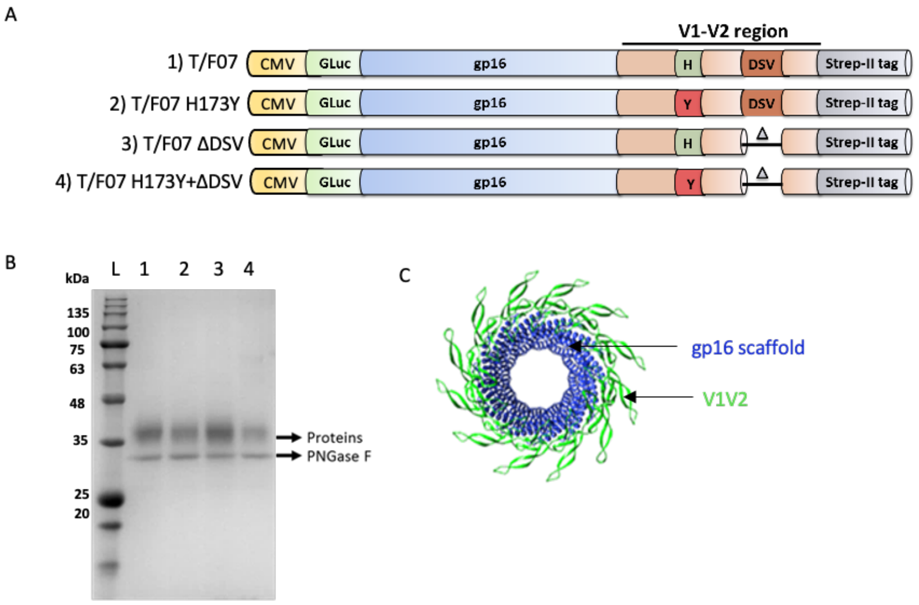


**Fig. S3. gp16-V1V2 construct design and purification. (A)** gp16-V1V2 constructs corresponding to the T/F07 virus sequence and the V1V2-specific mutations accumulated until 24-weeks of infection in participant 7. The V1V2 sequence (light orange) amplified from the T/F07 Env (gp160) sequence was fused in-frame to the C-terminus of gp16 scaffold (blue). Each construct was cloned under the control of the CMV promoter and contained an N-terminal signal peptide (GLuc) for secretion of these recombinant proteins into the medium, and a C-terminal Twin Strep-tag II (gray) for affinity purification. The 24-week V2 mutations were introduced in the C β-strand (H173Y, red) and the hypervariable V2 loop (3 residue deletion-ΔDSV) indicated by a small triangle in the parental T/F07 construct to generate single and double mutants. **(B)** SDS-PAGE profile of gp16-V1V2 variant scaffolds of T/F07 expressed in HEK293S (GnTi) cells and purified through StrepTactin affinity chromatography. These recombinant glycoproteins were deglycosylated by PNGase F (band labeled on the gel) to obtain sharper bands on the gel for the purpose of quantification. **(C)** Dodecameric model of gp16-V1V2 showing gp16-scaffold in blue and fused V1V2 domain in bright green.

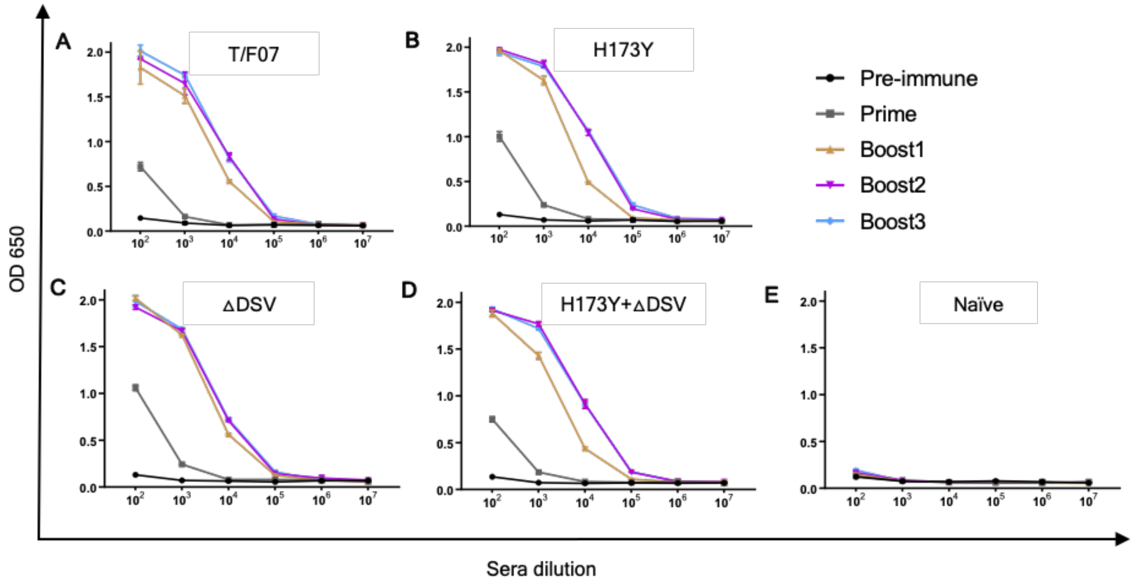


**Fig. S4. V1V2-specific antibody titers. (A-E)** Comparative titers of V1V2 antibodies ranging from the first (prime) to the last immunization (Boost 3) in mice groups immunized with gp16- T/F07 (A) H173Y (B) ΔDSV (C) H173Y+ΔDSV (D), and Naïve group (no antigen) (E) are shown. Respective pre-immune sera (collected before first immunization) from each group were also used as negative control. The antibody titers were determined by ELISA. Respective purified recombinant soluble gp140-T/F07, -H173Y, -ΔDSV and -H173Y+ΔDSV Env glycoproteins with matching V1V2 region were used as coating antigens (1 μg/ml). Triplicate absorbance (OD 650 nm) readings are used to generate binding curves for each sample. A color-coded key is provided on the top-right corner for each curve.

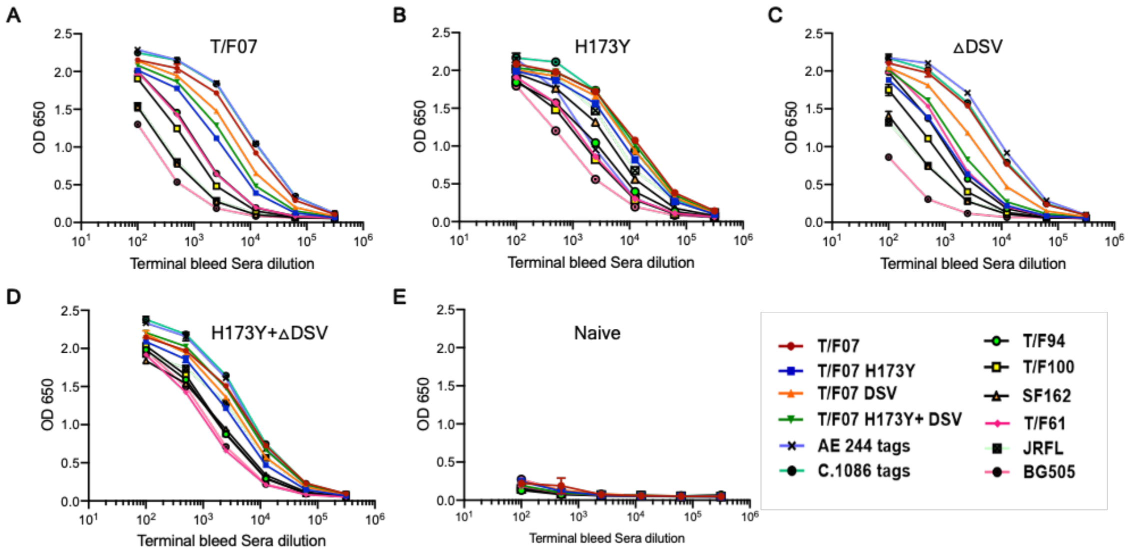


**Fig. S5. (A-E)** Breadth was evaluated by assessing binding with a set of purified autologous and heterologous Env proteins (soluble gp140s and V1V2 tags). Respective antigen binding curves are plotted against the serial dilutions of the terminal bleed sera from each group immunized with T/F07 (A), H173Y (B), ΔDSV (C), H173Y+ΔDSV (D), and PBS (Naïve, negative control) (E). Binding curves are color-coded with respect to the antigen coated as shown at the bottom right of the graph. The binding was determined through ELISA and triplicate absorbance (OD 650 nm) readings were used to generate the binding curves.

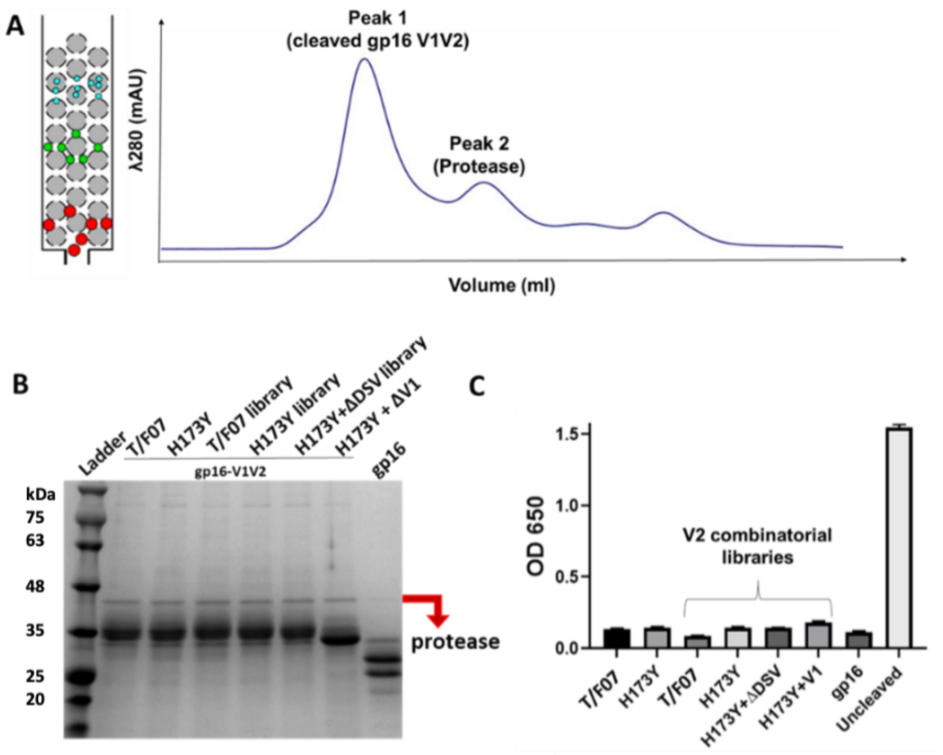


**Fig. S6. Purification of tag-free V1V2 immunogens. (A)** Size exclusion chromatography (SEC) fractionation profile of cleaved gp16-V1V2 proteins (peak 1) depicting separation of HRV3C protease (peak 2). Elution volume is plotted on the x-axis while the y-axis shows UV absorbance of the fractions. **(B)** Reducing SDS-PAGE profile of purified and concentrated gp16-V1V2 immunogens. Presence of very small fraction of the protease (marked by a red arrow) was detected. Appearance of smeary pattern or doublet bands are due to the presence of glycoforms. **(C)** ELISA showed no significant detection of α-StrepTag response in the final preparation of twin StrepTag cleaved immunogens. Uncleaved (with twin-strep tag) gp16-V1V2 was used a positive control.

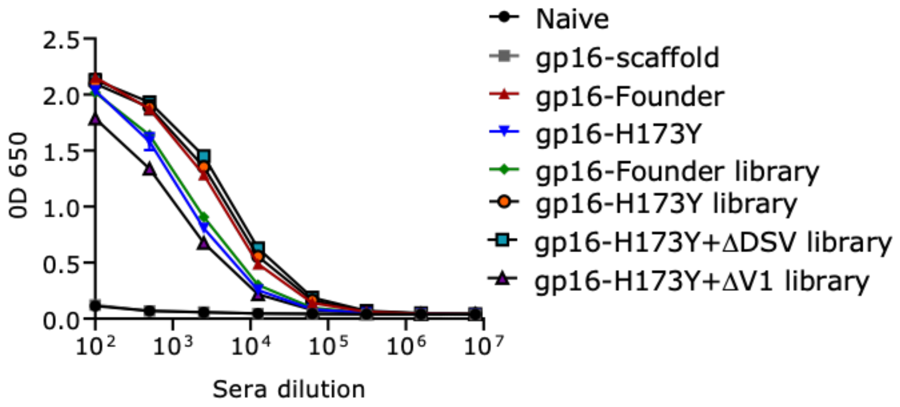


**Fig. S7. V1V2-specific binding responses in the terminal bleed sera.** V1V2-antibodies were detected in mice groups immunized with buffer (no antigen, Naïve group), gp16-scaffold only (no V1V2 control), gp16-V1V2- T/F07, H173Y, T/F07 library, H173Y library, H173Y+ΔDSV library and H173Y+ΔV1 library. A color-coded key is provided on the right side of the graph for each binding curve. Both naïve and gp16-scaffold only groups showed no non-specific reactivity towards the coating antigen. The antibody titers are determined through ELISA. Respective purified recombinant soluble gp140-T/F07, -H173Y, -ΔDSV, -H173Y+ΔDSV and –H173Y+ΔV1 Env glycoproteins were used as coating antigens (1 µg/ml) matching the V1V2 region (parental template mutations for combinatorial libraries). Triplicate absorbance (OD 650 nm) readings are used to generate the binding curves.

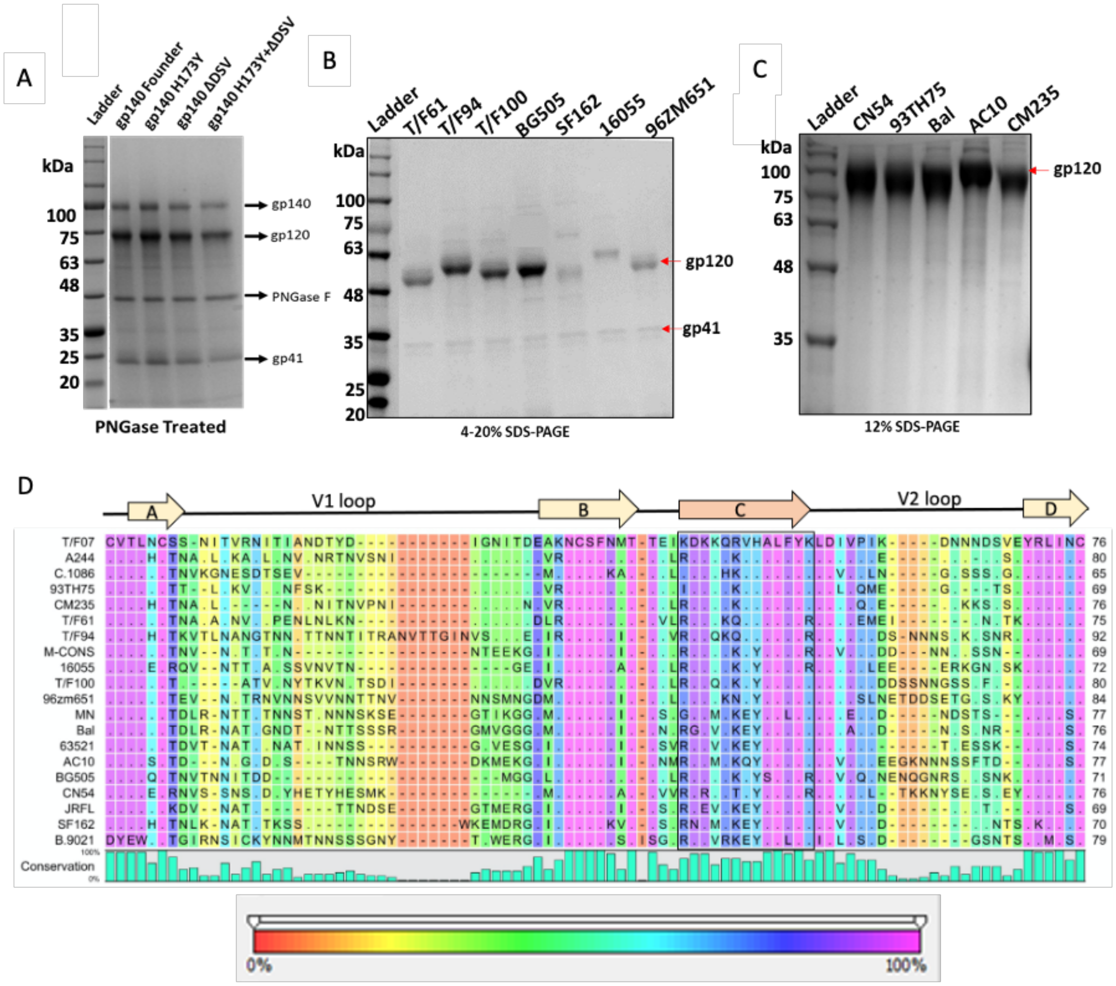


**Fig. S8. Recombinant construction and purification of diverse HIV-1 Env proteins to determine cross-reactive responses and breadth. (A-C)** Reducing SDS-PAGE profile of GnTi-expressed recombinant His-tagged Env proteins, gp140-T/F07 (Founder) and its V2 mutants (A); gp140s (cleaved into gp120 and gp41 subunits) (B); and gp120s (C) of different HIV-1 subtypes used as heterologous Env antigens. **(D)** V1V2 sequences of the diverse HIV-1 subtypes included in the heterologous Env protein library used to determine breadth. Degree of conservation (0-100%) at each residue position is depicted graphically at the bottom of the alignment. Variability in the V1V2 region of the chosen Env antigens is shown with background color gradient (red to pink) showing conservation on a scale of 0-100%.

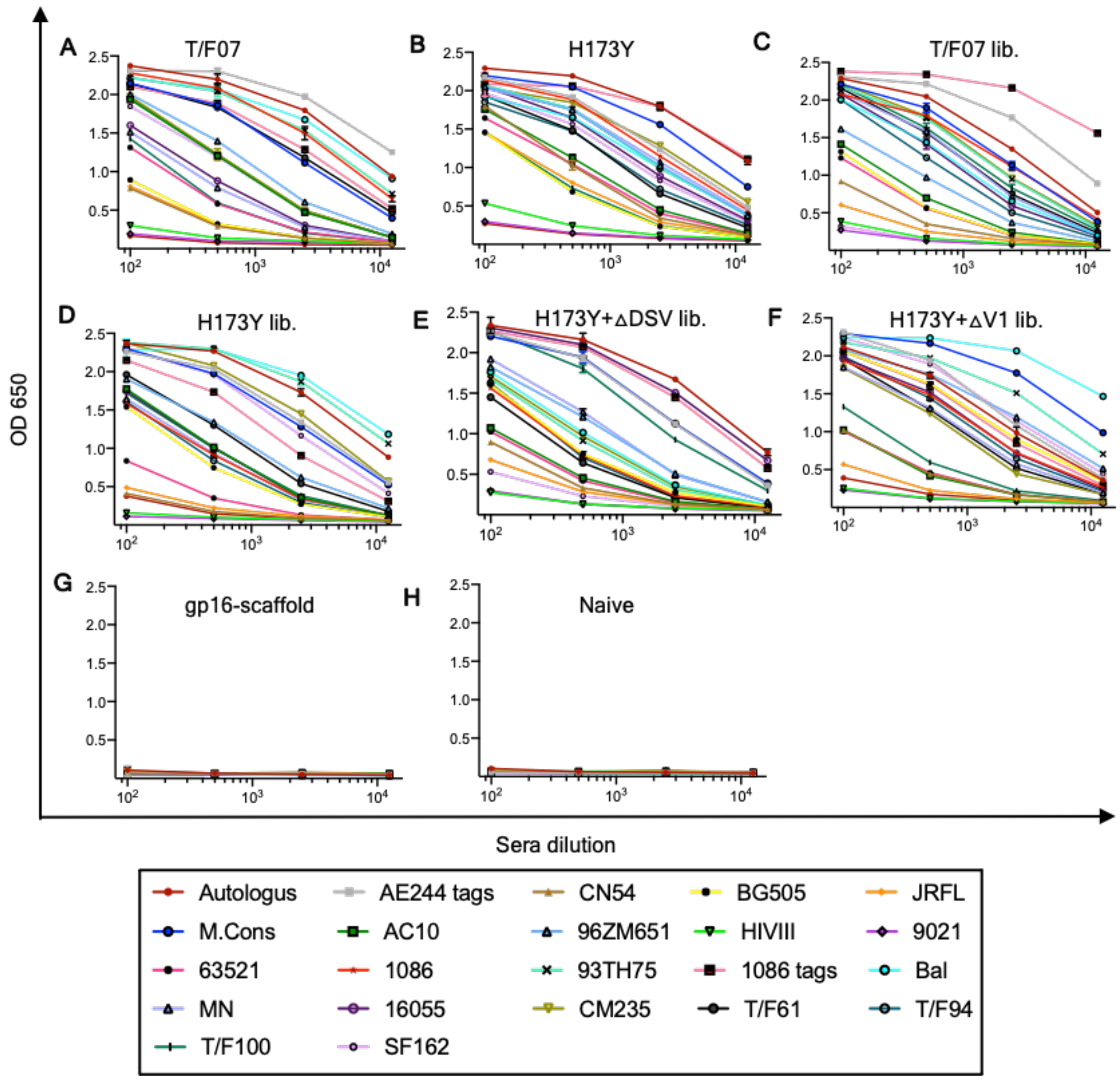


**Fig. S9. Breadth analysis of V2 combinatorial library immunogens using heterologous Env antigen library. (A-H)** ELISA generated binding curves showing the reactivity of sera of mice groups immunized with T/F07 (A) H173Y (B), and combinatorial V2 libraries (lib.); (C) T/F07 (D) H173Y (E) H173Y+ΔDSV (F) H173Y+ΔV1 groups (G). gp16-scaffold (H) and Naïve (I) groups sera were used as negative controls. The experiments were performed with 5-fold serially diluted pooled sera from each group in triplicates. Curves are color-coded for the respective antigen as shown in the box at the bottom.

**Table S1. List of reagents obtained from the NIH Reagent Program**

| Catalog # | Description |
|---|---|
| 4961 | HIV-1 BaL gp120 recombinant protein |
| 7749 | HIV-1 CN54 gp120 recombinant protein |
| 10080 | HIV-1 96ZM651 gp120 recombinant protein |
| 11556 | HIV-1 JR-CSF Fc-gp120 recombinant protein |
| 11784 | HIV-1 IIIB gp120 recombinant protein |
| 12063 | HIV-1 UG037 gp140 recombinant protein |
| 12064 | HIV-1 CN54 gp140 recombinant protein |
| 12569 | AE.A244 D11 gp120 recombinant protein |
| 12570 | B.MN D11 gp120 recombinant protein |
| 12571 | B.9021 D11gp120 recombinant protein |
| 12572 | B.6240 gp140C recombinant protein |
| 12574 | B.63521 D11 gp120 mutC recombinant protein |
| 12576 | M.CON-S D11 gp120 recombinant protein |
| 12581 | C.1086 gp140C recombinant protein |
| 13055 | HIV-1 AC10.29 gp120 Avi His recombinant protein |
| 13342 | HIV-1 93TH975 gp120 recombinant protein |
| 12567 | HIV-1 Env V1V2 recombinant protein (AE.A244 V1V2 tags) |
| 12568 | HIV-1 Env V1V2 recombinant protein (C.1086 V1V2 tags) |
| 8660 | HIV-1 96ZM651.8 gp140 optimized expression vector |
| 12806 | HIV-1 CM235 gp120 expression vector |
| 12957 | HIV-1 AC10.29 gp120 Avi His optimized expression vector |
| 13348 | HIV-1 BaL gp120 His expression vector |
| 13349 | HIV-1 93TH975 gp120 His expression vector |
| 13350 | HIV-1 CN54 gp120 His expression vector |
| 12551 | CH59 mAb |
| 12550 | CH58 mAb |
